# Acute Psychological Stress Triggers Circulating Cell-Free Mitochondrial DNA

**DOI:** 10.1101/405886

**Authors:** Caroline Trumpff, Anna L. Marsland, Carla Basualto-Alarcón, James L. Martin, Judith E. Carroll, Gabriel Sturm, Amy E. Vincent, Eugene V. Mosharov, Zhenglong Gu, Brett A. Kaufman, Martin Picard

## Abstract

Intrinsic biological mechanisms transduce psychological stress into physiological adaptation, but the role of mitochondria and mitochondrial DNA (mtDNA) in this process has not been defined in humans. Here, we show that similar to physical injury, psychological stress triggers elevation in circulating cell-free mtDNA (ccf-mtDNA). Healthy midlife adults exposed on two separate occasions to a brief psychological challenge exhibit a 2-3-fold increase in ccf-mtDNA, with no change in nuclear DNA levels, establishing the magnitude and specificity to ccf-mtDNA. In cell-based studies, we show that glucocorticoid signaling – a consequence of psychological stress in humans – is sufficient to induce mtDNA extrusion in a time frame consistent with human psychophysiology. Collectively, these findings provide the first evidence that psychological stress induces ccf-mtDNA and implicate glucocorticoid signaling as a trigger for ccf-mtDNA release. Further work is needed to examine the functional significance of psychological stress-induced ccf-mtDNA as a mitokine in humans.

## Introduction

Human health and disease risk is influenced by psychosocial factors transduced into adaptive biological changes via mechanisms that remain incompletely resolved ^1^. In response to perceived threat, humans and other mammals generate an integrated physiological response (the “fight-or-flight response”) involving the activation of multiple physiological systems. Every aspect of the stress response entails increased energy demand and thus necessarily engages mitochondrial energy production and signaling ^2^. The stress response is believed to have evolved to promote adaptation and increase the probability of survival ^3^; however, chronic activation of stress reactivity systems is associated with increased disease risk ^1,4^. Even brief exposure to a psychological stressor (i.e., an imagined threat) is sufficient to alter gene expression and elevate systemic markers of inflammation ^5,6^, reflecting the existence of intrinsic brain-body processes that transduce psychological stress into biological changes.

Recent animal studies suggest that chronic stress adversely influences multiple aspects of mitochondrial function and structural integrity ^7-9^ (reviewed in ^10^). Outside of the nucleus, mitochondria are the only organelle to contain their own genome – the mitochondrial DNA (mtDNA). Although the circular mtDNA is normally sequestered inside mitochondria, after physical stressors, such as trauma, infection, or strenuous exercise, mtDNA molecules are found in the circulation as circulating cell-free mitochondrial DNA (ccf-mtDNA) ^11-14^. Owing to its bacterial origin, ccf-mtDNA is immunogenic and triggers inflammation ^12,15^. During cell death, mtDNA is also actively released into the cytosol by selective molecular permeabilization of the mitochondrial membranes ^16^. Furthermore, ccf-mtDNA is also actively released by human lymphocytes and triggers immune activation ^17^, demonstrating that specific mechanisms exist to regulate mitochondrial genome release. Thus, given the ability of mitochondria-derived ccf-mtDNA to trigger inflammation, and that mitochondria are target of stress, these findings suggest that mitochondria could play a signaling role in response to threat, contributing to stress-induced immune activation and possibly other aspects of the stress response.

In addition to its elevation in injury and severe health conditions, higher ccf-mtDNA levels have been found in suicide attempters ^18^ and patients with major depressive disorder ^19^, representing cross-sectional evidence for a possible link between psychological states and ccf-mtDNA. Activation of the hypothalamic-pituitary-adrenal (HPA)-axis and the peripheral release of glucocorticoids is a primary neuroendocrine mediator of physiological responses to psychological stress ^20^. Interestingly, alterations of the HPA axis may also be implicated in the regulation of ccf-mtDNA levels in humans ^18^ and mtDNA gene expression in animals ^21^. However, whether psychological stress (in the absence of physical injury) or glucocorticoid signaling induce ccf-mtDNA release has not been established.

Accordingly, we investigated whether an acute psychosocial stress known to elicit the coordinated physiological stress response ^6,22,23^ is sufficient to affect ccf-mtDNA levels in humans. We sampled blood at three time points to examine dynamic changes in ccf-mtDNA levels in response to a social evaluative stressor, with a repeated challenge on a second visit one month later. To ascertain whether changes in ccf-mtDNA were due to non-specific release of bulk cellular material, we also assessed levels of circulating DNA from the nucleus (nDNA). Furthermore, given the evidence for sex differences in neuroendocrine response to acute stress ^24-26^ and in mitochondrial biology ^27^, we also explored sex differences in ccf-mtDNA levels. Finally, we used time lapse imaging in living human cells *in vitro* to test if glucocorticoid signaling is sufficient to trigger mtDNA extrusion. Our results show that exposure to a brief psychological stressor is sufficient to trigger a robust and specific increase in ccf-mtDNA, stronger in men than women, without a parallel increase in nDNA. In addition, glucocorticoid stimulation was sufficient to induce mtDNA extrusion from mitochondria. Overall, these findings implicate mitochondria and mtDNA signaling in the acute physiological response to psychological stress in humans.

## Results

Serum levels of two mtDNA (*mt-ND1*: mtDNA^1^, and *mt-CYTB*: mtDNA^2^) and two nDNA (*B2m*: nDNA^1^, and *Gusb*: nDNA^2^) amplicons were measured by qPCR (**Fig. 1*A,B***). This dual-amplicon approach insures that results are invariant to sequence differences that may exist between individuals, and differentiates between mitochondrial and nuclear genome release. A total of 50 healthy, midlife individuals (20 women, 30 men; mean age = 50 years, range: 41-58, 88% Caucasian) were studied. Serum was collected at three time points: i) *pre*: before the social evaluative stress test; ii) *post*: immediately after the stress test; and iii) *+30 min*: 30 minutes after the end of the stressor (**Fig. 1*C***). A validation visit took place one month later when all measures were repeated on the same participants. The detailed study design is illustrated in *Supplemental Information (SI)* (**Fig. S1)**.

**Fig. 1.**
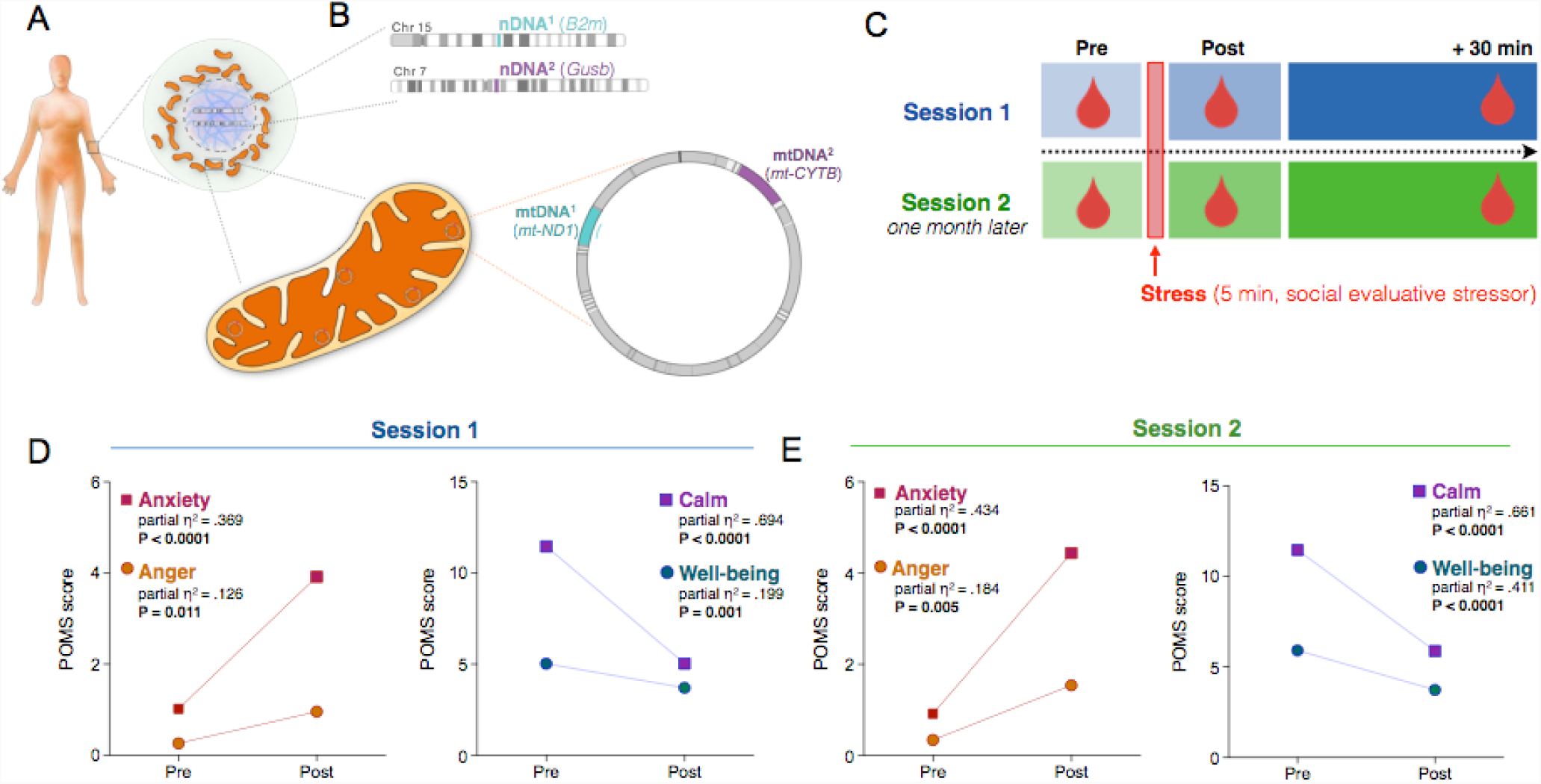
Study design to assess levels of circulating cell-free mitochondrial (ccf-mtDNA) and nuclear DNA (ccf-nDNA) in response to induced psychological stress. (**A**) Human cells contain nuclear DNA in the nucleus (purple) and cytoplasmic mitochondria (orange) with multiple copies of their own genome, the mtDNA. (**B**) Two different mtDNA amplicons, mtDNA^1^ (*mt-ND1*) and mtDNA^2^ (*mt-CYTB*), and two different nDNA amplicons nDNA^1^ (*B2m*) and nDNA^2^ (*Gusb*), were amplified from circulating cell-free serum using quantitative PCR. (**C**) Schematic of the experimental study design with three serial blood draws (pre, post, +30 min). A subsequent validation Session 2 was conducted one month later. (**D**) Stress-induced elevation of negative (left) and decrease in positive (right) items pre- and post-stress at Session 1 and (**E**) at Session 2, confirming the successful manipulation of psychological states. Paired two-tailed student T-tests, n = 49.

### ccf-mtDNA shows substantial inter- and intra-variability over one month

There were substantial inter-individual differences in baseline (pre-task) ccf-mtDNA^1^ (*mt-ND1*) and ccf-mtDNA^2^ (*mt-CYTB*) levels between sessions 1 and 2 (one month apart). The average between-person coefficient of variation (C.V.) for both mtDNA amplicons at both sessions was 98% (***SI*, Fig. S2**). In comparison, the C.V. for intra-individual (within-person) variation over a month was 33%. Moreover, for each participant baseline mtDNA levels were only moderately correlated between sessions 1 and 2 (***SI***, **Fig. S2*E,F***), indicating that baseline ccf-mtDNA shares both characteristics of a stable individual “trait,” but also exhibit substantial variation representing a variable “state” characteristic. For the nuclear genome, the between-person C.V. for both nDNA amplicons at both session was 69%, the within-person variability was 37%, and session 1 and 2 baseline ccf-nDNA values were not correlated (***SI*, Fig. S2**).

### Acute psychological stress increases ccf-mtDNA levels

Participants were exposed to a 5-minute psychological stress task involving the preparation (2 min) and delivery (3 min) of a speech to defend themselves against a false accusation (***SI*, Fig. S1**). To validate the effectiveness of this task as an experimental stressor, we examined indices of individual psychological states (i.e., mood) from *pre*- to *post*-task. Consistent with our previous report ^23^, at both sessions, the stressor caused a significant increase in negative mood (anxiety, anger) and decrease in positive mood (calm, well-being) (**Fig. 1*D-E***), demonstrating successful manipulation of the psychological state.

In response to this brief psychological challenge, ccf-mtDNA levels increased from *pre* to *+30 min* post-stress, exhibiting a large effect size (mtDNA^1^, 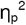= .569, *P*<0.0001, **Fig. 2*A*,** mtDNA^2^,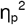 = 0.600 - 0.556, *P*<0.0001, ***SI*, Fig. S3*A***). The majority of participants (93%) showed an increase in both ccf-mtDNA^1^ and ccf-mtDNA^2^ after 30 minutes. Interestingly, ccf-mtDNA levels did not change significantly immediately after the stress, between *pre* and *post* samples, indicating some delay in this response.

**Fig. 2.**
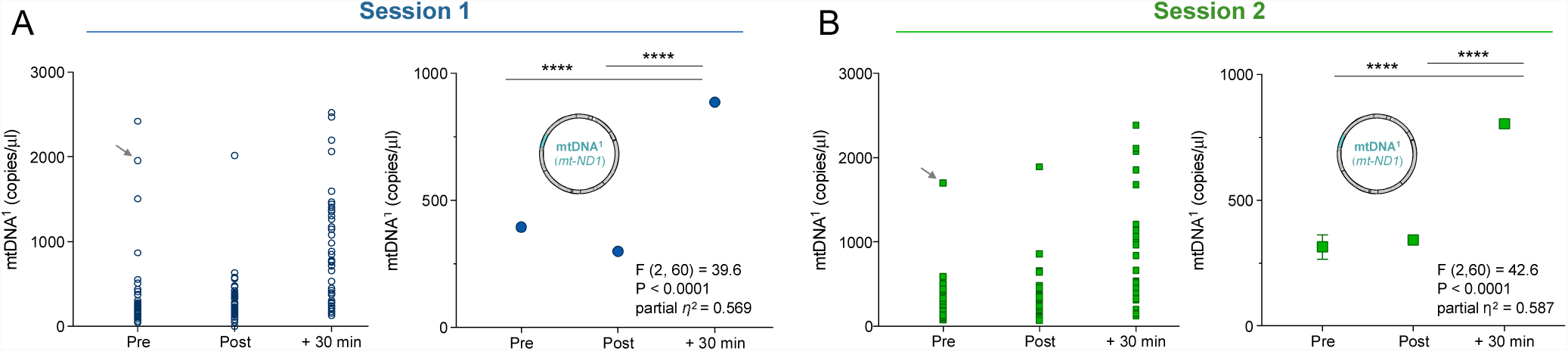
Psychological stress increases serum circulating cell-free mitochondrial DNA. (**A**) Individual (left) and group mean (right) ccf-mtDNA^1^ (*mt-ND1*) responses to acute psychological stress at session 1. (**B**) Validation of the results in (A) in a repeat session 2 one month later. Arrow: a participant with high mtDNA^1^ baseline values. Data are means and SEM. Repeated measure ANOVA on log transformed data and Least Significant Difference (LSD) pairwise comparisons, n = 31 per session, * P < 0.05, ** P < 0.01, *** P < 0.001, **** P < 0.0001.

The significant release of mtDNA at *+30 min* post stress was replicated at a second visit, one month later. Again, individuals showed a large increase in the levels of both ccf-mtDNA amplicons, with 94% of participants showing an increase in both ccf-mtDNA^1^ and ccf-mtDNA^2^. The effect size for session 2 was comparable to session 1 (mtDNA^1^, 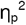 = .587 *P*<0.0001, **Fig. 2*B*** mtDNA^2^, 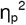 = .556, *P*<0.0001, ***SI*, Fig. S3*B***). The replication of this stress-induced mtDNA release confirmed that similar to physical injury and sepsis ^11^, psychosocial stress is sufficient to trigger robust elevations in serum ccf-mtDNA levels in human subjects.

### Stress selectively increases ccf-mtDNA

We next examined whether the increased ccf-mtDNA was specific to mitochondria or reflected a general increase in circulating total cellular genomic material. Thus, the same serum samples were analyzed for nuclear genome content. In contrast to ccf-mtDNA, circulating levels of nDNA^1^ (*B2m*) or nDNA^2^ (*Gusb)* did not increase in response to stress. In fact, both nuclear sequences tended to decrease at *+30 min* (nDNA^1^: 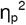 = .162 *P*<0.01, **Fig. 3*A***; nDNA^2^: 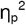 = .174 *P*<0.05, ***SI*, Fig. S4*A***). Because the mitochondrial and nuclear genomes were quantified in duplex reactions (i.e. in the same reaction), this result cannot be due to a sampling error. We also confirmed the ccf-nDNA profile in the validation session one month later, where ccf-nDNA^1^ and nDNA^2^ were unchanged (nDNA^1^, 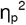 = 0.051, *P* = 0.20, **Fig. 3*B***; nDNA^2^, 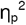 = 0.068, *P* = 0.132, ***SI*, Fig. S4*B***).

**Fig. 3.**
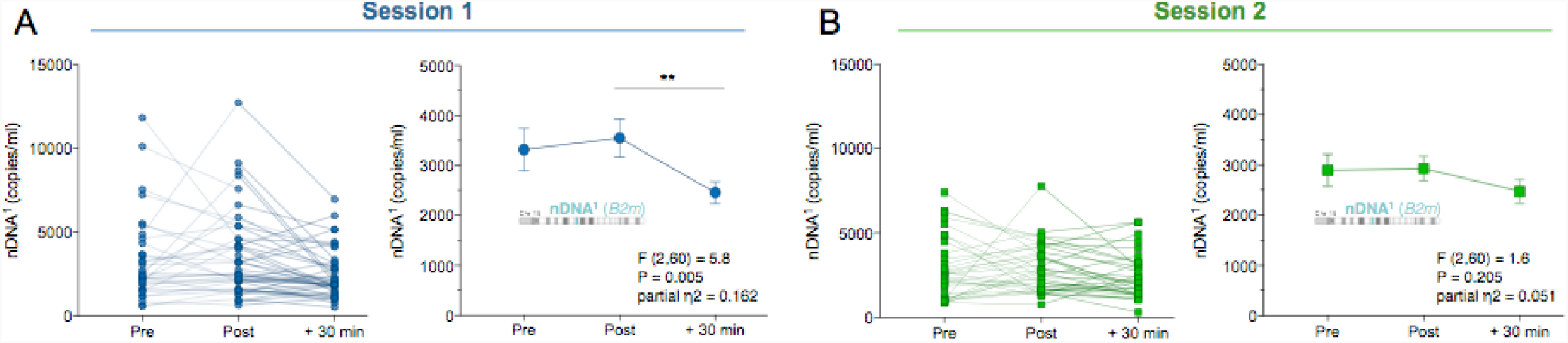
Psychological stress does not increase serum circulating cell-free nuclear DNA. (**A**) Responses to acute psychological stress at session 1 for the nuclear DNA (ccf-nDNA^1^, *B2m*) and (**B**) at the validation session 2. Data are means and SEM. Repeated measure ANOVA on log transformed data and Least Significant Difference (LSD) pairwise comparisons, n = 31 per session, * P < 0.05, ** P < 0.01, *** P < 0.001, **** P < 0.0001.

Furthermore, the ratio of the number of circulating mitochondrial to nuclear genomes (mtDNA/nDNA) also illustrates the selective increase of mtDNA over nDNA after stress. The baseline mtDNA/nDNA ratio was ~140 for DNA^1^ and ~63 for DNA^2^ (***SI*, Fig. S5*A-D***). Relative to *pre* ratio, the mtDNA^1^/nDNA^1^ ratio at *+30 min* rose by > 2-fold during both first and second visits (*P*<0.0001, ***SI*, Fig. S5*A,B***). The direction and magnitude of effect was confirmed with the second set of amplicons mtDNA^2^/nDNA^2^ (***SI*, Fig. S5*C,D***). Together, these data show that psychological stress selectively increases circulating cell-free mitochondrial but not nuclear DNA, and that their circulating levels are likely modulated by different biological mechanisms.

### Stress-induced ccf-mtDNA elevation differs by sex

Next, we conducted secondary analyses to investigate whether sex moderated the magnitude of stress-induced ccf-mtDNA responses. The complete results of sex-stratified analyses of the levels of ccf-mtDNA in response to stress are presented in **SI Fig. S6**. In a model that included both sessions, a significant sex by time interaction was found for mtDNA^1^ (F (2,105) = 4.60, P < 0.05, 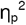 = 0.195) and mtDNA^2^ (F (2,120) = 5.68, P < 0.01, 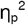 = 0.087) (**Fig. 4*A,B***), with higher + 30 min ccf-mtDNA levels observed in men, suggesting that psychological stress-induced ccf-mtDNA release may differ by sex. The effect sizes also reveal consistently larger effects of stress on ccf-mtDNA for men than women (***SI*, Fig. S7**), particularly at session 2. In comparison, the effects sizes for nDNA were substantially smaller than for mtDNA and showed no evidence of a sex difference at either session (***SI*, Fig. S7*B***). This finding further reinforces the notion that psychological stress-induced DNA release is specific to the mitochondrial genome.

**Fig. 4.**
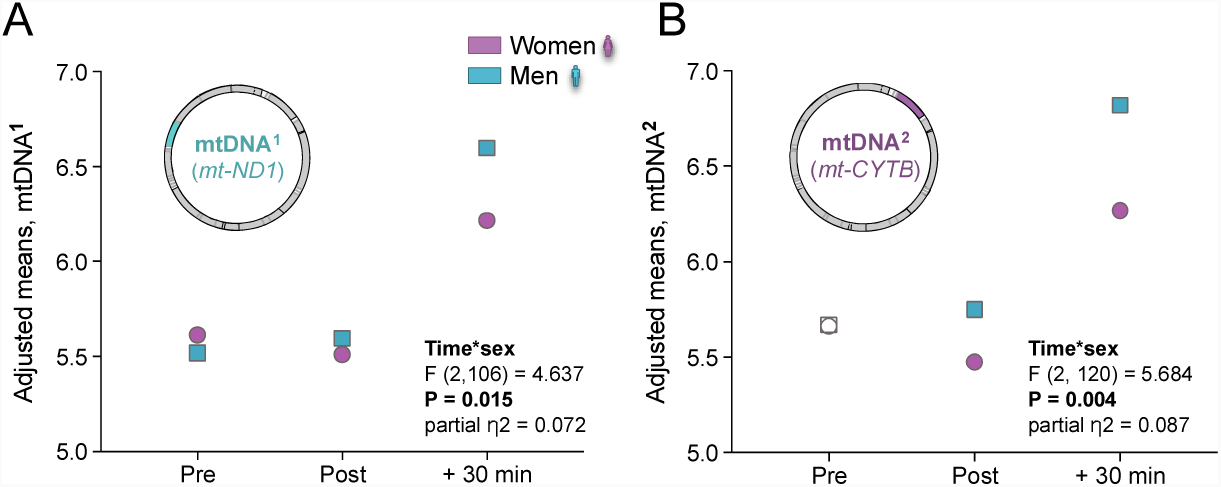
Stress by sex interaction on the levels of circulating cell-free mtDNA. (**A**) Adjusted means for ccf-mtDNA^**^1^**^ responses to acute psychological stress. (**B**) Same as in (A) for ccf-mtDNA^**^2^**^. Repeated measure ANOVA on log transformed data, n = 62 across both sessions.

### Glucocorticoid signaling triggers mtDNA extrusion in living cells

Finally, to explore a potential cellular mechanism for stress-induced ccf-mtDNA levels, we tested glucocorticoid signaling as a potential trigger of mtDNA extrusion in human cells. We stimulated primary human fibroblast *in vitro* for up to an hour with the glucocorticoid hormone mimetic dexamethasone (DEX, 100nM) (**Fig. 5A**) and concurrently quantified the intracellular distribution of the glucocorticoid receptor (GR). As expected, DEX stimulation caused a rapid redistribution of GR from the cytoplasm to the nucleus peaking at 30 min, demonstrating the activation of glucocorticoid signaling in this system (**Fig. 5B-C**).

**Fig. 5.**
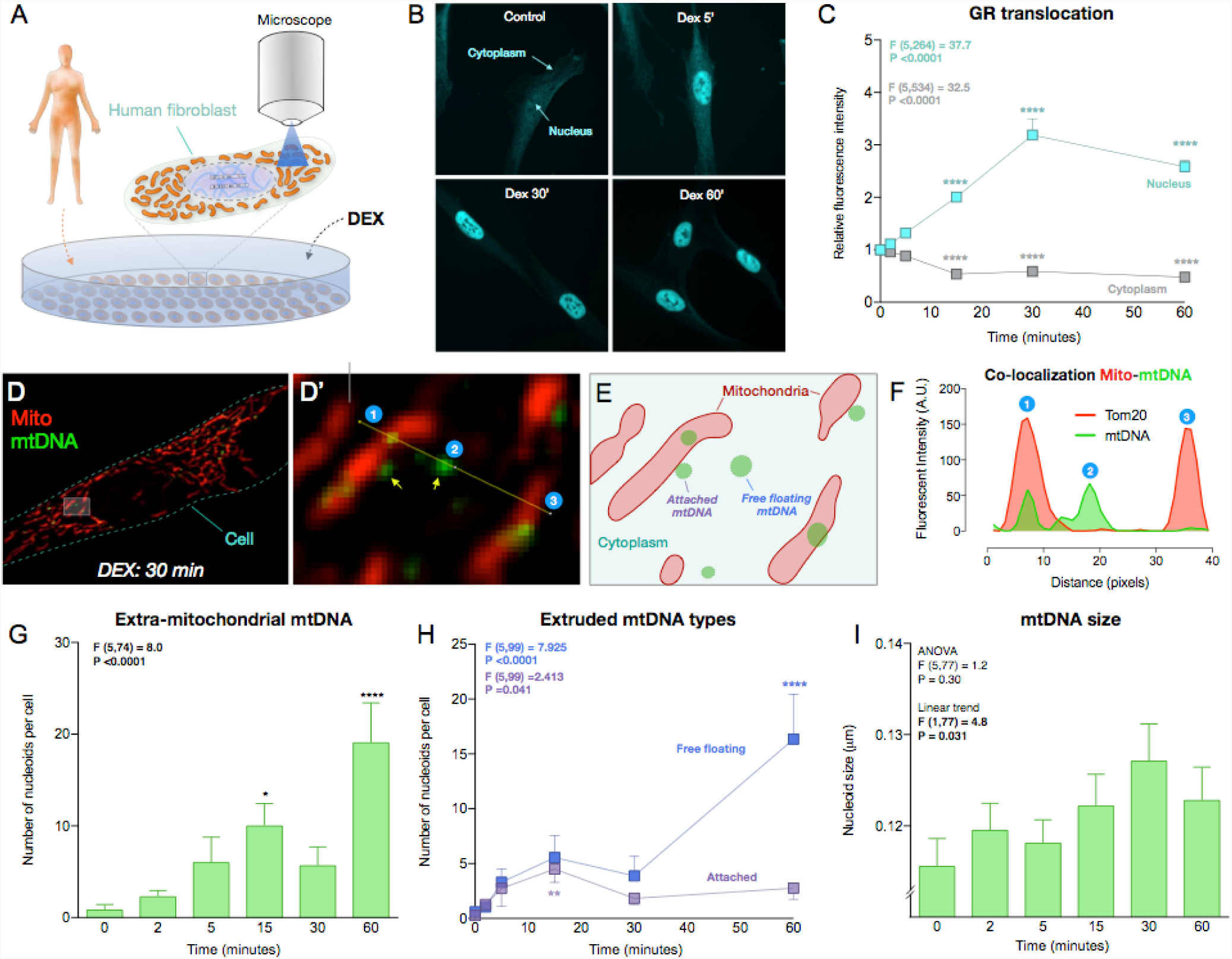
Glucocorticoid stimulation causes mtDNA extrusion in human cells. (**A**) Schematic of the experimental design. Primary male human fibroblasts were acutely exposed to 100nM dexamethasone (DEX) for 0, 2, 5, 15, 30 and 60 minutes, and mitochondria and mtDNA visualized by immunofluorescence. (**B**) Representative pictures of human fibroblasts after DEX treatment analyzed for GR subcellular localization. (**C**) Glucocorticoid receptor (GR) fluorescence was quantified in both the nucleus and the cytoplasm following DEX stimulation, which induced a marked translocation of GR to the nucleus. One Way ANOVA and Least Significant Difference (LSD) pairwise comparisons. (**D**) A human fibroblast exposed to DEX stimulation for 30 minutes, and dual-labeled by immunofluorescence for a ubiquitous marker of mitochondria (Tom20) and mtDNA. (**D’**) Higher magnification of boxed area in (D). (**E**) Cartoon of the mitochondria and nucleoids observed in (D’). Note the two different forms of extruded mtDNA: in contact to the surface of mitochondria (attached), or free floating in the cytoplasm (free floating). (**F**) Line profile intensity quantification for Tom20 and mtDNA along the yellow line in D’ showing: 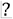 A nucleoid inside the mitochondrion; 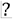 A nucleoid extruded into the cytoplasm (no mito-Tom20 staining); and 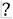 A mitochondrion without mtDNA. (**G**) Time-course of the number of extruded mtDNA nucleoids after acute DEX stimulation. One Way ANOVA and Least Significant Difference (LSD) pairwise comparisons, n = 82 cells. (**H**) Time-course of sub-types of mtDNA nucleoids: attached and free floating mtDNA. One Way ANOVA and LSD pairwise comparisons, n = 105 cells. (**I**) Size of extruded mtDNA (averages per cell). One Way ANOVA and Post hoc test for linear trend, n = 105 cells. Data are means ± SEM. Pairwise comparisons presented are relative to DEX 0 minute. * P < 0.05, ** P < 0.01, *** P < 0.001, **** P < 0.0001.

During DEX stimulation, we simultaneously tracked mitochondria and mtDNA. Under normal conditions, >99.5% of mtDNA appear as punctate structures, termed nucleoids, inside mitochondria.Strikingly, acute glucocorticoid stimulation increased the number of mtDNA nucleoids located outside mitochondria as early as 15 min (**Fig. 5D-G**). Extruded mtDNA were categorized as “mitochondrion- attached” or “free-floating,” presumably representing the early and late stages of mtDNA extrusion, respectively. Accordingly, the number of attached nucleoids showed a gradual increase peaking at 15 minutes, and free-floating mtDNA increased and peaked at 60 minutes (**Fig. 5H**). This process was associated with increased mtDNA nucleoid size (+10.4% at 30 min) (post hoc test for linear trend F (1,77) = 4.8, *P* < 0.05), consistent with the release and “decondensation” of mtDNA known to occur during other forms of cellular stress (**Fig. 5I**) ^28,29^.

## Discussion

The organism’s ability to adapt to stress depends on the concerted action of molecular factors secreted within minutes to hours, enabling the transfer of information across physiological system, which ultimately promotes survival of the organism. The present study demonstrates that a brief psychological stressor is sufficient to cause a robust and rapid increase in serum ccf-mtDNA, implicating the mitochondrial genome as a stress-inducible cytokine, or “mitokine.” On both occasions of testing, the results showed a selective increase in ccf-mtDNA levels, but not ccf-nDNA, within 30 minutes. Consistent with preclinical studies showing deleterious effects of acute psychological stress on mitochondrial structure and function ^10^ and previous work showing elevated ccf-mtDNA in suicide attempters ^18^, these experimental results identify ccf-mtDNA as a biological marker of mitochondrial stress secondary to acute psychological challenge in humans, and identify glucocorticoid signaling as a potential cellular trigger of this effect.

Clinically, increases in ccf-mtDNA have been ascribed to tissue damage and higher levels predict mortality ^11^. In a prospective study of 443 critically ill patients in the intensive care unit, elevated ccf-mtDNA levels were a strong predictor of mortality, associated with a 7-fold increased risk of death within the next month ^13^. Levels of ccf-mtDNA also appear to increase with age. In a study of 831 healthy individuals, compared to children and young adults aged 1-41 years, older adults aged 50-89 showed 3.8 fold higher ccf-mtDNA levels, and those >90 years showed a 7.0-fold elevation ^15^. Together, these findings raise the possibility that levels of ccf-mtDNA increase with age and are functionally significant, predicting increased disease and mortality risk ^11^. Thus, further understanding of mechanisms that regulate ccf-mtDNA levels is warranted.

In relation to psychological stress, our findings support initial cross-sectional evidence in two psychiatric populations – non-violent suicide attempters and patients with major depressive disorder – both of whom show elevated levels of plasma ccf-mtDNA when compared with matched controls ^18,19^. In our study, the elevation *pre*/*post*- to *+30 min* across two sessions yielded Cohen’s *d* values ranging from 0.85-1.23, demonstrating a robust and rapid effect of psychological stress on serum ccf-mtDNA. Other than mtDNA, mitochondrial proteins such as heat shock protein 60 (Hsp60), cytochrome c, and prohibitins have also been detected in the circulation of healthy individuals ^30-32^. In one study, psychological distress, high levels of job demand, and low levels of emotional support were found to be associated with circulating plasma mitochondrial Hsp60 levels ^31^. Our findings extend these data by showing that serum ccf-mtDNA is acutely responsive to the psychological state, and thus should be considered as a dynamic stress-inducible marker.

The current study also broadens prior assumptions that ccf-mtDNA is a consequence of physical injury to raise the possibility that ccf-mtDNA may also contribute to the increased risk of morbidity and mortality associated with chronic psychological stress. However, the full downstream physiological effects of acute increases in ccf-mtDNA remain to be determined. One possibility is that ccf-mtDNA contributes to the delayed effects of psychological stress on inflammation and metabolism ^11,33^. Mitochondria are the only organelle with their own DNA, which is recognized as bacterial-like and activates pattern-recognition receptors both intracellularly and extracellularly ^34^. In animals, injection of mtDNA in the blood triggers systemic inflammation in a toll-like receptor 9-dependent manner ^12^. Furthermore, in humans, inflammatory diseases have been associated with elevated ccf-mtDNA ^35^, and the anti-inflammatory mediator acetylcholine may prevent mtDNA release from mitochondria ^36^. Consistent with this idea, older individuals with elevated circulating levels of ccf-mtDNA tend to have higher levels of proinflammatory cytokines IL-6 and TNF-α, and mtDNA functions as an adjuvant that potentiates the production of TNF-α in isolated monocytes ^15^. Having established the rapid inducibility of ccf-mtDNA, further research is thus warranted to determine if the increase in ccf-mtDNA secondary to acute psychological stress is necessary and sufficient to trigger stress-induced inflammation in humans.

Indeed, psychological stress is known to evoke systemic inflammation ^6,37^. In humans, stress-evoked inflammatory responses peak around 90-120 minutes following the onset of stress ^6^. Thus, the present study shows that the elevation in serum ccf-mtDNA at 30 minutes precedes the established time frame for the inflammatory response, which would be consistent with the immunogenic nature of the mtDNA. We previously showed in this cohort that higher negative emotional responses to the stressor were positively associated with IL-6 responses from baseline to 30 minutes after stress ^23^. But since we observed a ccf-mtDNA increase at the same timepoint, and could not assess later time points in this cohort, the current study does not allow us to test whether ccf-mtDNA increase contributed to subsequent inflammation.

Moreover, in line with some, but not all, previous reports suggesting sex differences in the magnitude of neuroendocrine responses to social-evaluative stress ^24-26^, we observed a greater stress-related increase in ccf-mtDNA in men compared to women. While further work is needed to explain these findings, it should be noted that several aspects of mitochondria biology, including respiratory capacity, sensitivity to permeability transition, and reactive oxygen species production have been shown to differ by sex, with increased vulnerability generally observed in males ^27^. There are also significant metabolic differences between women and men ^38^ and some mitochondrial disorders show sexual dimorphism even at a young age before differences in sex hormones appear, with boys being more susceptible than girls ^39,40^. In relation to ccf-mtDNA, resolving these and other questions will require addressing certain limitations, including the need for denser and longer kinetic analyses of ccf-mtDNA in relation to neuroendocrine and inflammatory markers. In addition, future studies should distinguish between plasma and serum levels since ccf-mtDNA concentration measured in serum may be substantially higher due to additional release of cellular DNA during coagulation in serum ^41^.

Finally, it must be noted that the source of circulating DNA in humans, particularly in response to psychological stress, is not clear. The bulk release of cellular material as a result of cell death would result in an increase in both mitochondrial and nuclear genomes. In contrast, the current findings show a selective increase in ccf-mtDNA levels with no concomitant increase in nDNA. These dissociable responses to social-evaluative stress strongly argue against cellular death as the mechanism of stress-induced increases in ccf-mtDNA. Another possible mechanism involves the active release of mtDNA. Recent evidence describe molecular mechanisms of mtDNA release from the mitochondrial matrix to the cytoplasm ^16^ and from the cell to the circulation ^17^, providing a biological basis for rapid and selective extrusion of mtDNA. Our data on primary human fibroblasts showing that glucocorticoid signaling is sufficient to rapidly trigger extrusion of mtDNA outside of mitochondria suggests that canonical neuroendocrine stress mediators may represent a link between stressful experiences and mtDNA release. Additional work is required to establish, in different cell types, the mechanisms for stress-induced mtDNA extrusion.

## Conclusions

Overall, this experimental study demonstrates that psychological stress selectively increases serum ccf-mtDNA levels in humans. The current findings add to a growing literature on circulating DNAs ^11^, providing initial evidence that elevation of ccf-mtDNA occurs not only with physical injury, inflammatory diseases, critical illness, and aging, but also in response to acute psychological stress. Furthermore, this study provides evidence that glucocorticoid signaling induces mtDNA extrusion in human cells, suggesting a potential pathway linking subjective stressful psychological states to mitochondrial signaling. Overall, this work implicates mitochondrial adaptations and mitokine signaling in the fight-or-flight response, and possibly in human stress pathophysiology.

## Materials and Methods

### Study cohort

Samples and data for the present study were obtained from the Vaccination and Immunity Project, a longitudinal study investigating the association of psychosocial, physiologic, and behavioral factors with antibody response to hepatitis B vaccination in a middle age adult population ^23,42^. A total of 50 participants (30 men, 20 women, 88% Caucasian) aged between 41-58 years were included in the present study and 32 participants (64%) completed both visits.

Participants were non-smokers, in good general health (no history of myocardial infarction, asthma, cancer treatment in the past year, psychotic illness, or other systemic diseases affecting the immune system) free from medication interfering with nervous, endocrine, and immune system (excepted for oral contraceptives) for the 3 months prior to the beginning of the study. Pregnant and lactating women were ineligible. Participants were free of symptoms of infection or antibiotic use for the 2 weeks prior to the laboratory visits. Informed consent was obtained in compliance with guidelines of the University of Pittsburgh Institutional Review Board.

### Experimental stress procedure

The detailed study design is illustrated in Supplemental Information (SI) (**Fig. S1*A***) and has been previously reported ^23,42^. Prior to receiving the hepatitis B vaccination series, participants attended two laboratory sessions scheduled 1 month apart. On each occasion, participants abstained from caffeine, food, and beverage (excepted water) for 12 h, non-prescription medications and physical activity for 24 h and alcohol beverages for 48 h before coming into the laboratory between 7:00 and 9:00 AM. On arrival, an intravenous catheter was inserted into the antecubital fossa of one arm for collection of blood samples. Participants then rested quietly for a 30-minute adaptation period and a *pre*-task sample of blood was drawn and mood states were assessed using the Profile of Mood States questionnaire (POMS) ^43,44^. Participants then performed a simulated public speaking task, consisting of 2 minutes of preparation for a speech defending themselves against an alleged transgression (shoplifting or traffic violation) followed by 3 minutes of videotaped speech delivery (see 23,42). We have shown previously that this task elicits reproducible cardiovascular and immune stress responses ^23,42,45,46^. Mood states were collected and *post*-task blood samples were drawn immediately following completion of the task and again after the subject had rested quietly for 30 minutes (*+30 min*). The procedure was the same at the second laboratory visit except subjects were told that their “performance on the first speech task was slightly below average when compared with other participants’ speeches,” and participants were asked to “try to be more persuasive when delivering this speech.”

### Serum DNA extraction and quantification

Blood samples were allowed to clot, centrifuged at 1,000 × g for 10 minutes, and the serum was frozen at -80.°C until further processing (**Fig. S1*B***). Each sample was thawed, visually inspected, and subjected to an additional centrifugation at 2,000 × g for 5 minutes in 1.5ml microfuge tubes to prevent potential cellular contamination prior to DNA extraction.

There was no relationship between sample color (yellow, pink due to erythrocyte lysis, or opacity) and the amount of DNA detected. Similar to the previously reported fraction of mtDNA molecules that are freely circulating and those enclosed in small vesicles (~19%) ^47^, removal of membrane-bound fractions by centrifugation of plasma at 10,000 × g and 18,000 x g reduced circulating mtDNA levels by ~31% in our samples. All analyses shown are therefore representative of both free DNA and small membrane-bound pools of circulating mtDNA.

Total nucleic acids were extracted from serum by proteinase K and ethanol precipitation as previously described ^48^, and circulating levels of mtDNA measured by duplex quantitative real-time PCR (qPCR) with Taqman chemistry ^49^. Linearity of assays was confirmed using titration of mixed total DNA from 30 human placentas, which was used to calculate relative abundance of individual samples, and read counts validated by digital PCR. Two different reactions were run in parallel for each sample, with each measure performed in triplicates, including a standard curve for each reaction plate. Reactions consisted in both a mtDNA and nDNA amplicons: *mt-ND1* and *B2m* for mtDNA^1^ and nDNA^1^, respectively; and *mt-CYTB* and *Gusb* for mtDNA^2^ and nDNA^2^, respectively (see *SI Table 1* for details of primers/probes). Two different amplicons for both the mitochondrial and nuclear genomes were used to ensure that mtDNA and nDNA quantifications are invariant of potential inter-individual sequence variation. Both mtDNA^1^ and mtDNA^2^ yielded highly correlated results (**Fig. S1*D***).

### Dexamethasone stimulation of primary human fibroblast

Primary human male fibroblasts (hFB1m) obtained from a healthy donor (passages 9-13) were seeded on cover slips and grown in DMEM supplemented with non-essential amino acids and physiological glucose concentration (5.5mM). After 24 hours, cells were acutely exposed to the synthetic glucocorticoid receptor agonist dexamethasone (DEX, 100 nM) for 0, 2, 5, 15, 30 and 60 minutes. Triple immunofluorescence microscopy was used to localize the glucocorticoid receptor (GR), mitochondria, and mtDNA.

#### Immunofluorescence and image analysis

Following DEX stimulation, cells were fixed with 4% paraformaldehyde (PFA) for 15 minutes. Cells were washed 3 times with PBS and then permeabilized with 90% Methanol (diluted in PBS) for 20 minutes at -30°C. Cells were washed with PBS three times and blocked with 10% Normal Goat Serum (NGS) for 60 minutes at room temperature (RT). Primary antibody incubation was as follows: Tom20 (Santa Cruz (F10); sc-17764), mouse monoclonal IgG2a, 1:100 in 2% NGS; Glucocorticoid receptor (GR) (Santa Cruz (G-5); sc-393232, which recognizes aa. 121-420), mouse monoclonal IgG2b, 1:100 in 2% NGS; Anti-DNA antibody (American Research Product (AC-30-10); #03-61014), mouse monoclonal IgM, 1:100 in 2% NGS. Coverslips were incubated at 4°C, overnight. For secondary antibody incubation cells were first washed 3 times with PBS and then incubated in darkness for 60 minutes at RT, as follows: Mouse IgG2a-594, 1:500 in 2% NGS; Mouse IgG2b-488, 1:500 in 2% NGS; Mouse IgM-AMCA, 1:500 in 2% NGS. Cells were washed 3 times with PBS and mounted with ProLong Gold (Thermo Fisher Scientific # P10144). Images were taken on an Olympus IX81 inverted fluorescence microscope equipped with a digitized stage (ProScan; Prior Scientific), a 63x/1.35 oil objective (Olympus, MA), a corresponding fluorescence filter set and a 2.0 neutral density filter using a CoolSNAP HQ camera (Roper Scientific/Photometrics, AZ) and MetaMorph software (Molecular Devices, CA).

For GR translocation analyses, at least five images were analyzed per time point, and experiments repeated three times. For each cell analyzed, regions of interest (ROIs) were averaged for the cytoplasm (six ROIs), nucleus (three), and background (three). For analyses of extruded mtDNA located outside mitochondria, a mask using the mitochondrial channel (Tom20) was used to remove mtDNA nucleoids inside mitochondria, and nucleoid number and size were automatically determined. Each extruded mtDNA particle was analyzed manually and classified as being in immediate contact with a mitochondrion (attached) or without any overlapping pixel with mitochondria (free floating). Nucleoid size was also measured for all nucleoids at each time point and expressed as μm^2^. A line scan was performed to assess co-localization of mitochondria and mtDNA nucleoids (see Figure 5). All images were analyzed for densitometry or colocalization using Image J ^50^, and all data is the average of three separate experiments.

### Data analysis

Statistical analyses were performed using SAS 9.3, SPSS 23, and Prism 7.0 (Graphpad). mtDNA and nDNA data obtained from the human study were natural log transformed before analyses or analyzed using non-parametric tests. mtDNA/nDNA ratio was computed to estimate the relative abundance of mitochondrial and nuclear DNA levels. Linear regression was used to compare nDNA and mtDNA abundance between sessions and to investigate association between mtDNA and nDNA at *pre*, *post* and *+30 min*. To evaluate main effects of the stressor on mtDNA and nDNA concentration, repeated-measures analyses of variance (ANOVAs) were conducted, followed by Least Significant Difference (LSD) pairwise comparisons when indicated. Measures of effect size were calculated using partial eta-squared (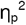). For analyses where the assumption of sphericity (tested using Mauchly’s test) was not met, the Greenhouse-Geisser correction was applied. In a model that included both sessions, we tested for a sex x time interaction using repeated-measures analyses of covariance (ANCOVAs). Sex differences were further explored by stratifying analysis. For cell culture experiments, difference in mtDNA nucleoids and GR characteristics between the DEX conditions (0, 2, 5, 15, 30, 60 minutes) were assessed using One Way ANOVA, post hoc test for linear trend (when non-significant), and LSD pairwise comparisons. Two-tailed statistical significance was accepted as *P*<0.05.

### Data availability

The data supporting the findings of the present report are available from the corresponding authors upon request.

## Author contributions

ALM, JEC and MP designed the study. ALM and JEC conducted the original study and provided samples. AV and MP processed samples. JLM and BAK performed ccf-DNA measurements and analyses. CBA performed in vitro experiments with GS. CT performed statistical analysis. CT and MP drafted the manuscript. All authors contributed to the final version of the manuscript.

